# Loss of COX4i1 leads to combined respiratory chain deficiency and impaired mitochondrial proteosynthesis

**DOI:** 10.1101/2020.01.31.925420

**Authors:** Čunátová Kristýna, Pajuelo Reguera David, Vrbacký Marek, Fernández-Vizarra Erika, Ding Shujing, Fearnley Ian, Zeviani Massimo, Houštěk Josef, Mráček Tomáš, Pecina Petr

## Abstract

Oxidative phosphorylation (OXPHOS) system localized in the inner mitochondrial membrane secures production of the majority of ATP in mammalian organisms. Individual OXPHOS complexes were shown to form supramolecular assemblies termed supercomplexes. It has been repeatedly shown that complexes are not linked only by their function but also by interdependence of individual complex biogenesis or maintenance. For instance, cytochrome *c* oxidase (cIV, COX) or cytochrome *bc1* complex (cIII) deficiencies affect the level of fully assembled NADH dehydrogenase (cI) in monomer as well as within supercomplexes. It was hypothesized that cI is affected at the level of enzyme assembly as well as at the level of cI stability and maintenance. However, the true nature of interdependency between cI and cIV is not fully understood yet. We used HEK293 cellular model with complete knockout of COX4 subunit, which serves as an ideal system to study interdependency of cI and cIV, as early phases of cIV assembly process are disrupted. Total absence of cIV was accompanied by profound deficiency of cI, documented by selective decrease in cI subunits amount and significantly reduced amount of assembled cI. Supercomplexes assembled from cI, cIII and cIV were missing in COX4dKO due to loss of cIV and decrease in cI amount. Pulse-chase metabolic labelling of mtDNA-encoded proteins uncovered decrease of cIV and cI subunits translation. Moreover, partial impairment of mitochondrial proteosynthesis correlated with decreased level of mitochondrial ribosomal proteins. In addition, complexome profiling approach uncovered accumulation of cI assembly intermediates indicating that cI biogenesis was affected rather than stability. We propose that impairment of mitochondrial proteosynthesis caused by cIV deficiency represents one of the mechanisms which may couple biogenesis of cI and cIV.

## INTRODUCTION

Energetic needs of mammalian organisms fully depend on ATP produced by oxidative phosphorylation (OXPHOS), a major metabolic pathway localized in the inner mitochondrial membrane. OXPHOS complexes are encoded by both mitochondrial (mtDNA) and nuclear (nDNA) DNA. Complex I (NADH dehydrogenase, cI) contains seven out of 45 subunits encoded by mtDNA, complex II (succinate dehydrogenase, cII) contains 4 nuclear-encoded subunits, complex III (cytochrome bc1 complex, cIII) contains one of 10 subunits encoded by mtDNA, and complex IV (cytochrome *c* oxidase, cIV, COX) contains three out of 14 subunits encoded by mtDNA. Regulation of OXPHOS capacity may therefore occur at various levels of mtDNA expression, including mRNA maturation, mRNA stability, translational coordination, ribosomal biogenesis, and translation itself [1–5]. Expression and posttranslational modifications of individual subunits in cytosol and mitochondria need to be coordinated by signalling crosstalk between nucleus and mitochondria. Also, for proper OXPHOS function, many enzyme cofactors are needed and their homeostasis and efficient incorporation into apoenzyme are important [6]. Individual subunits and cofactors must assemble to produce the mature and functional enzymes, with a myriad of so-called ‘assembly factors’ involved. In addition, respiratory chain complexes form supramolecular assemblies termed as supercomplexes, adding yet another level of complexity in OXPHOS biogenesis. The most abundant supercomplex in mammalian mitochondria is the respirasome, composed of one copy of complex I, dimer of complex III and one copy of COX (I_1_III_2_IV_1_) [7]. Its structure has been recently solved with overall resolution at 5.8 Å [8] or 5.4 Å [9].

Biogenesis of mammalian COX is a complicated and highly regulated process that was first described as sequential incorporation of COX subunits through four distinct assembly intermediates S1 – S4, reflecting rate-limiting steps in enzyme biogenesis [10]. The COX assembly mechanism is not spontaneous, because numerous accessory nuclear DNA encoded proteins are needed to build the holoenzyme. Their function is required for all steps of the process and most of them are evolutionary conserved from lower to higher eukaryotes [11,12]. Currently, COX assembly is considered to be modular, with independent maturation of three mtDNA-encoded subunits accompanied by addition of specific sets of nuclear-encoded subunits [4,13]. COX1 maturation occurs within the MITRAC complexes (mitochondrial translation regulation assembly intermediates of COX), responsible for integration of mitochondrially encoded and newly imported nuclear encoded proteins with the help of TIM21 protein [14]. The function of early-MITRAC complex lies in regulation of COX1 translation by coordinating the interaction of COX1 with specific assembly proteins (COX14, MITRAC12 – hCOA3, MITRAC7) before proceeding to the further steps of COX assembly process. Insertion of heme *a* and *a3* into COX1 occurs in late-MITRAC complexes, where SURF1 is recruited along with enzymes responsible for heme a synthesis [15], before the addition of nuclear-encoded subunits COX4 and COX5a. COX2 module then associates with the COX1-COX4-COX5a intermediate upon its copper metallization by the specific SCO1/SCO2 copper chaperones [16]. Addition of COX3 module and the majority of nuclear-encoded subunits finalizes COX assembly [4].

It has been repeatedly shown that deficiency in one of the complexes may in turn affect other complexes. Therefore, complexes are linked not only by their function, but also by interdependence of individual complex formation or maintenance. Notably, deficiency of cIII has been repeatedly shown to induce decrease of cI content and activity. While it has been traditionally interpreted as cI destabilization in the absence of its supercomplex partner [17–22], some studies convincingly show that the interdependency is associated with impairment in cI maturation, which can be at least partially attributed to essential association of both these complexes within the respirasome during their assembly [23,24].

Less frequently, it was shown that cIV deficiency affects level of assembled cI both non-associated and within supercomplexes. Interestingly, this phenomenon seems to occur only in rare cases with complete absence of cIV caused by defect in early phases of COX1 maturation/assembly [25,26]. These studies indicate that cI is affected at the level of enzyme assembly. Complex I is the largest OXPHOS complex, which in mammals consists of 45 subunits, including 14 core subunits homologous to prokaryotic cI components, and numerous accessory subunits that are indispensable for enzyme assembly and stabilization [5]. The biogenesis of the complex is characterized by pre-assembly of individual modules that reflect enzyme functional units, assisted by specific assembly factors [4]. The membrane arm containing the mtDNA encoded subunits ND1-ND6 first associates with the ubiquinone (coQ)-binding module. The assembly is completed by attachment of the soluble N-module responsible for electron transfer between NADH and CoQ. In this regard, functioning of some of the cI assembly factors may be affected by altered redox status of mitochondria following loss of complex IV, such as proteins (e.g. AIF) involved in oxidative Mia40 pathway-dependent oxidative folding of intermembrane space residing components of complex I, or factors responsible for synthesis and delivery of FeS clusters into the matrix arm of cI. Other case of cI-cIV interdependency reported that cIV loss due to nonsense mutations in COX1 gene is associated with elevated activity of mitochondrial m-AAA protease AFG3L2, and therefore impairs the stability of complex I rather than its biogenesis [27]. Importantly, numerous cases of milder cIV assembly defects with at least minimal residual portion of assembled cIV display no effect on cI [28].

Here, we report that complete removal of COX4 subunit in HEK293 results in complete COX deficiency. Furthermore, the impairment of the earliest steps in COX assembly also causes profound deficiency of complex I, which is associated with decreased synthesis of mtDNA-encoded cI subunits as well as reduced content of mitochondrial ribosomal proteins. We propose that mitochondrial proteosynthesis attenuation may represent one of the mechanisms of cIV-cI biogenesis interdependency.

## MATERIALS AND METHODS

### Cell culture

HEK293 (human embryonic kidney) cells were cultivated under standard conditions (37 °C, 5% CO_2_ atmosphere), initially in high-glucose DMEM medium (Life Technologies, 31966-021) supplemented with 10% (v/v) FBS (Life Technologies, 10270-106), 20 mM HEPES and antibiotics (100 U/mL penicillin, 100 μg/mL streptomycin, Life Technologies, 15140-122). After construction of COX4 knock-out cellular models, cultivation was switched into more nutrient-rich DMEM/F-12 (Life Technologies, 31331-028), concentration of HEPES buffer was doubled to 40 mM, and additional nutrient supplement, 50 μM uridine, was used to facilitate growth of highly glycolytic knock-out cells.

### Preparation of HEK293 cellular knock-out models

COX4i1 and COX4i2 gene knock-out was introduced into HEK293 cells by CRISPR knock-out technology. Specifically, Cas9-D10A paired nickase with two chimeric RNA duplexes was used for its higher specificity for targeted sequence [29]. gRNAs for COX4i1 and COX4i2 were obtained from commercially available library (Sigma-Aldrich, USA) in the form of expression plasmid (U6-gRNA, expressed by U6 promoter), along with expression plasmid for Cas9-D10A nickase fused with GFP (Sigma-Aldrich, USA). For COX4i1 KO, HSL0001377215/ HSR0001377224 gRNA pair was used, and HSR0002687616 / HSL0002687619 gRNA pair was utilized for COX4i2 gene KO. HEK293 cells were transfected by relevant plasmids, incubated for 72 hours and then trypsinized, and diluted to concentration of 20 cells/mL. This suspension was aliquoted by 100 μL onto 96-well plates. Wells containing single cell colonies were identified and further cultivated. Confluent cells in 6-well plates were harvested and used for selection of mutant clones by sequencing and SDS-PAGE. Two clones of each COX4i1, COX4i2, and COX4i1/2 with full loss of respective protein on the background of nonsense mutation generating premature stop codons were chosen for the study.

### COX4 isoform overexpression

COX4i1 (CCDS10955.1) and COX4i2 (CCDS13187.1) isoforms of human COX4 subunit full length cDNAs including 3’ end sequence encoding FLAG tag (DYKDDDDK) were obtained as synthetized DNA strands with 5’ and 3’ overhangs containing restriction sites used for insertion into pcDNA3.1^+^ mammalian expression vector (ThermoFischer). COX4i1-FLAG or COX4i2-FLAG constructs were transfected into COX4i1/2 KO cells using Metafectene Pro (Biontex), stably-transfected cells were selected with 1 mg.mL^−1^ G418.

### SDS-PAGE

Tricine sodium dodecyl sulphate polyacrylamide gel electrophoresis (SDS-PAGE) was used for separation of proteins based on their different molecular weight under denaturing conditions [30]. SDS-PAGE samples were prepared from frozen aliquots of harvested cellular pellets in SLB buffer (sample lysis buffer; 2% (v/v) 2-mercaptoethanol, 4% (w/v) SDS, 50 mM Tris (pH 7,0), 10% (v/v) glycerol, 0,02% Coomassie Brilliant Blue R-250) to final concentration 2-5 mg protein/mL. Then, samples were sonicated and incubated for 20 min in 40 °C. Samples (10-50 μg of protein as indicated) were separated on 12 % gel using Mini-PROTEAN III apparatus (Bio-Rad, USA).

### Native electrophoresis

Blue-native (BNE-PAGE) electrophoresis [31] was used to separate native protein complexes. For protein complexes solubilization, mild non-ionic detergent digitonin, which prevents dissociation of supercomplexes, was used, thus OXPHOS complexes and their supercomplexes can be detected. Mitochondria from HEK293 were released by hypotonic shock and isolated using differential centrifugation as described [32] and then stored as dry pellets at −80°C. Isolated mitochondrial pellets were resuspended in solubilisation buffer A (50mM NaCl, 50mM imidazole/HCl, 2mM 6-aminocaproic acid, 1mM EDTA; pH 7,0) to reach final protein concentration 10 mg/mL. Digitonin (20% (w/v) stock solution) was added (6g detergent/ g of protein) to mitochondrial suspension, and solubilisation proceeded on ice for 10 min. Afterwards, samples were centrifuged at 30 000 g (20 min, 4 °C) and supernatant containing released membrane proteins was transferred into clean tube. Concentration of protein and sample volume were measured. Then, glycerol was added to final 5 % (v/v) concentration and Coomassie Brilliant Blue G-250 (CBB) was supplemented according to detergent/dye ratio 8:1 (w/w). Samples were separated on 5 – 16 % gradient polyacrylamide gel using Mini-PROTEAN III apparatus (Bio-Rad, USA).

### Western blot and immunodetection

Proteins separated by electrophoresis were transferred onto PVDF (polyvinylidene difluoride) membrane (Immobilon FL 0,45 μm, Millipore Corporation, USA) by semi-dry electroblotting using Transblot SD apparatus (Bio-Rad Laboratories, USA). After blotting, PVDF membrane was washed for 5 min in TBS (150 mM Tris-HCl, 10 mM NaCl; pH 7,5) and blocked in 5% (w/v) fat-free dry milk diluted in TBS for 1 hour. Then, membrane was washed 2×10 min in TBST (0,1% (v/v) detergent Tween-20 in TBS) and was prepared for immunodetection of proteins.

For immunodetection, membrane was incubated for 2 hours in primary antibody (diluted in TBST) at room temperature. Then, membrane was washed 3×10 min with TBST and secondary antibody (conjugated with fluorescence tag, diluted in TBST) incubation for 1 hour in dark followed. Afterwards, membrane was washed 3×10 min in TBST and 10 min in TBS. Primary and secondary antibodies used for immunodetection are mentioned in Supplementary table 1. Signal detection was performed using fluorescence scanner Odyssey (LI-COR Biosciences, USA). Signals detected by scanner were quantified by software AIDA Image Analyzer (Raytest Isotopenmessgeräte GmbH, Germany).

### Mass spectrometry – label free quantification (MS LFQ)

Biological duplicates (COX4i1 KO clones 1 and 2, COX4i2 KO clones 1 and 2, COX4i1/4i2 KO clones 1 and 2) were harvested and MS LFQ analysis was performed in technical duplicates as described [33]. Briefly, cell pellets (100 μg of protein) of wild-type HEK293 cell line, COX4i1, COX4i2 and COX4i1/4i2 KO clones were solubilized by sodium deoxycholate (final concentration 1% (w/v)), reduced with TCEP [tris(2-carboxyethyl)phosphine], alkylated with MMTS (S-methyl methanethiosulfonate), digested sequentially with Lys-C and trypsin and extracted with ethylacetate saturated with water as described. Samples were desalted on Empore C18 columns, dried in Speedvac and dissolved in 0.1% TFA + 2% acetonitrile. About 1 μg of peptide digests were separated on 50 cm C18 column using 2.5 h elution gradient and analyzed in a DDA mode on a Orbitrap Fusion Tribrid (Thermo Scientific) mass spectrometer. Resulting raw files were processed in MaxQuant (v. 1.5.3.28) with label-free quantification (LFQ) algorithm MaxLFQ. Downstream analysis and visualization was performed in Perseus (v. 1.5.3.1).

### SILAC/Complexome Profiling

Wild-type and COX4 dKO cell lines were cultivated in “heavy” (^15^N- and ^13^C-labeled Arg and Lys) and in “light” (^14^N and ^12^C Arg and Lys) DMEM (Thermo). Unlabeled (“light”) cell line was mixed equally with labeled (“heavy”) one and mitochondria-enriched fraction was isolated [34]. Mitochondrial pellets were solubilized by digitonin and Bis/Tris BN-PAGE was performed using 3-12% gradient gel. The gel was stained, fixed and cut to 64 slices. Slices were in-gel digested by trypsin and analyzed by tandem mass spectrometry [13,35]. For the protein identification, peptide quantification and plain text export, Proteome Discoverer software (Thermo) and Mascot peptide search engine (Matrix Science) were used. For further data evaluation, Python code was used to separate unlabeled and labeled peptides into two datasets and R code was used to determine the relative peptide intensity for each protein according to the most frequently occurring representative peptide in both unlabeled and labeled forms (for details see [13]). The relative peptide intensity profile of each protein was normalized to the slice with the highest intensity and average of reciprocal experiments was visualized in ‘heatmaps’ using Microsoft Excel and Prism 7 (GraphPad Software, La Jolla California USA).

### Pulse-chase analysis

To study proteosynthesis and turnover of mtDNA-encoded proteins, the method of *in-vivo* metabolic pulse-chase protein labelling by ^35^S-methionine + ^35^S-cysteine was employed essentially as described [36], using Express Protein Labelling Mix, [^35^S] (PerkinElmer, USA). Cells were labelled in DMEM-M-C with 100 μg/mL of anisomycin and the radioactive labelling mix with 100 μCi/mL 37 °C and 5% atmosphere for 3 hours. After 3 hours, 20 μL of the mixture of “cold” methionine and cysteine (non-labelled Met and Cys) was added into each well (final concentration 250 μM) and incubated at 37 °C and 5% CO2 atmosphere for 15 min. Then, cells were washed two-times with PBS containing 250 μM cold methionine and 250 μM cold cysteine. One aliquot of labelled cells was immediately harvested (“pulse” sample). Two other labelled cells aliquots (“chase” samples) were further incubated in complete non-labelling DMEM medium for 4 or 24 hours. Cellular pellets were later used for SDS-PAGE analysis using large format Hoefer™ SE 600 Chroma Vertical Electrophoresis System (Thermofisher, USA). 70 μg protein-aliquots of each sample were separated by 16 % gel. Following the electrophoretic run, gel was fixed in CBB staining solution (40 % (v/v) methanol, 8 % (v/v) acetic acid, 0.05 % (w/w) Coommasie Brilliant Blue R-250) and dried onto Whatmann CHR 3 mm paper. Radioactivity was detected by exposing the gel to Storage Phosphor Screen BAS-IP SR 2025 E for 14 days (GE Healthcare), which was then scanned by FX Molecular Imager (Bio-Rad Laboratories, USA). Signals detected by scanner were quantified by software AIDA Image Analyzer (Raytest Isotopenmessgeräte GmbH, Germany).

### Measurements of metabolic fluxes

Parallel measurement of mitochondrial respiration and glycolytic rate was performed using Seahorse Extracellular Flux (XF) Analyzer (Agilent Technologies, USA). One day prior to measurement, 3.105 cells were were seeded in poly-L-lysine coated wells of measuring plate. Four wells were always prepared without cells to serve for calibration of background signals. Microplate was then incubated overnight in thermostat. Hydration of XF24 Sensor Cartridge by XF Calibrant solution (pH 7,4; Agilent Technologies, USA) was also carried out overnight at 37 °C. Day after, microplate was washed with 1 mL of XF Assay Medium Modified DMEM containing 10 mM glucose as a substrate (pH 7,4, 37 °C), 500 μL of the same medium with 0.2 % (w/w) BSA was pipetted and microplate was incubated at 37 °C for 30 min. Meanwhile, XFe24 Sensor Cartridge was prepared by injection of substrates and inhibitors to record basal metabolic rate with glucose and after subsequent additions reaching final concentrations of 1 μM oligomycin (Oligo), 1 μM FCCP (FCCP), and inhibitor cocktail of 1 μM rotenone, 1 μg/ml antimycin A, and 100 mM 2-deoxyglucose (Rot+AA+2DG). For precise normalization of rates according to cell counts, immediately after Seahorse run, cell nuclei were stained by Hoechst dye (final concentration 5 μg/mL, diluted in FluoroBrite DMEM Media). Images of whole wells were acquired by Cytation 3 Cell Imaging Reader (BioTek, USA) and analysed using software Gen5 (BioTek) to obtain cell counts for each well.

## RESULTS

### Decreased steady-state content of COX and cI subunits in cells lacking COX4i1

The steady-state content of five representative COX subunits (COX4i1, COX1, COX2, COX5a, and Cox6c) was examined by denaturing SDS-PAGE followed by Western blot analysis and immunodetection in two representative clones of each COX4i2 KO, COX4i1 KO and COX4i1/4i2 KO clones in comparison to wildtype (wt) HEK293. Representative WB images are shown in Fig. 1A, where COX4i1, COX2 were undetectable, COX5a and COX6c only barely detectable, and COX1 significantly decreased in COX4i1 KO and double COX4i1/4i2 KO (Fig. 1A, S1 A,B). In contrast, COX4i2 KO cells showed similar pattern as the wt. The functionality and specificity of the antibodies used to detect Cox4i1 or Cox4i2 was confirmed using positive control samples from COX4i1/4i2 KO transfected with constructs carrying cDNA of the corresponding genes (Fig. 1B). Densitometric quantification revealed that relative levels of studied COX subunits in COX4i2 KO clones ranged between 55 – 120 % of controls with significant decrease of Cox2 content, when data were normalized to actin levels (Fig. S1 A). However, upon normalization to citrate synthase levels (marker of mitochondrial mass), COX subunits content was unaffected (Figure S.1 B). These data suggest that COX4i2 KO have mildly decreased amount of mitochondria rather than selectively decreased COX content. (Fig. S1 A,B).

**Figure 1:**
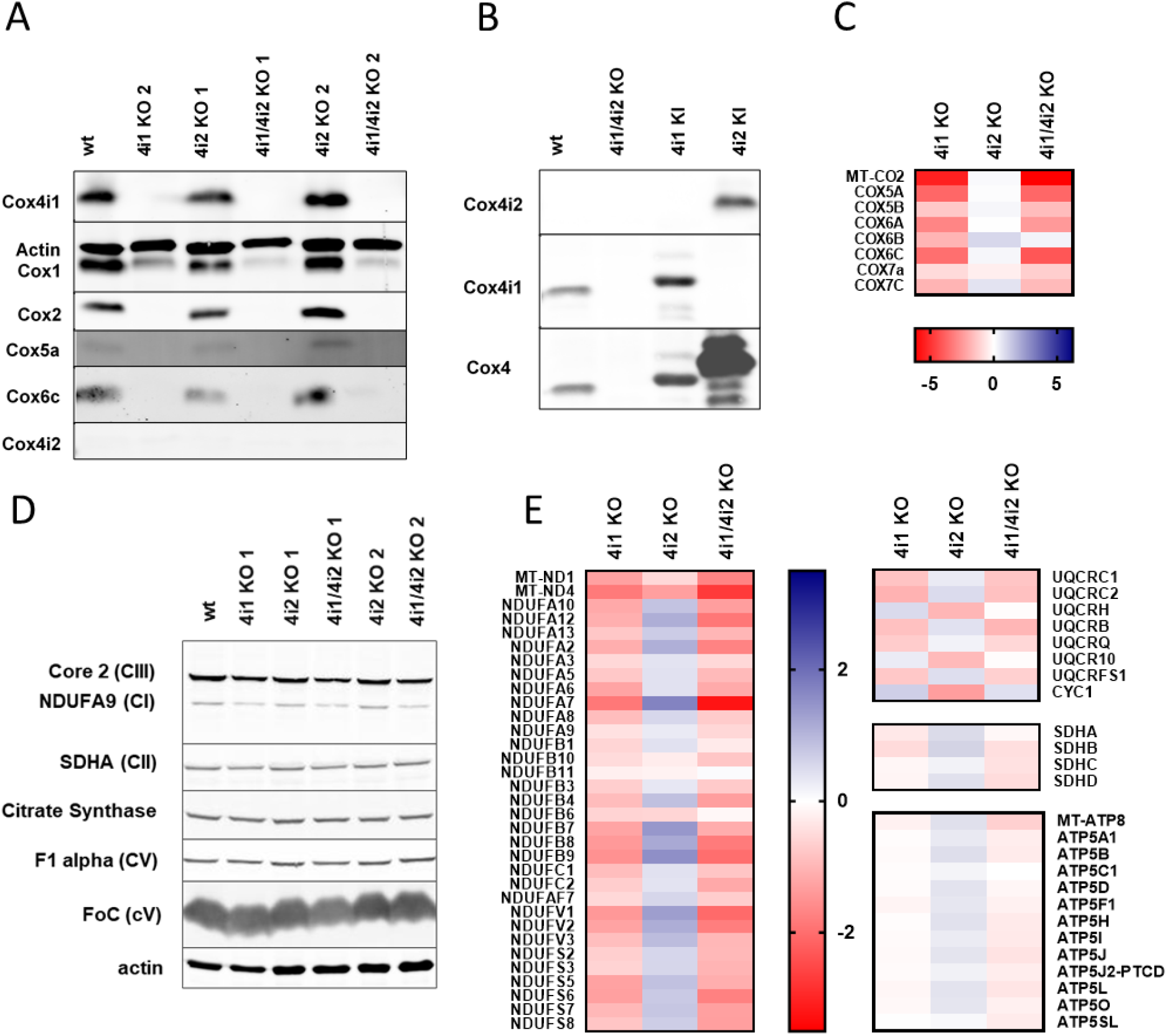
**a) Representative Western blot analysis of COX subunits in KO clones.** Whole cell lysates (30 μg of protein) of HEK293 (wt) COX4i1 KO, COX4i2 KO and COX4i1/4i2 KO clones were subjected to SDS-PAGE and Western blotting. Actin antibody was used as a loading control. **b) COX4i2 antibody testing.** Whole cell lysates (30 μg of protein) of COX4i1 KO, COX4i2 KO and COX4i1/4i2 KO clones were subjected to SDS-PAGE and Western blotting. HEK293 cells (wt) do not show detectable levels COX4i2 protein. On the contrary, COX4i1 isoform was detected by both, monoclonal (COX4i1) and polyclonal (non-specific COX4) antibodies. COX4i2 protein was detected in positive control, COX4i2 knock-in. **c) Proteomic analysis of relative content of COX subunits using mass spectrometry label-free quantification (MS LFQ).** Figure shows decreased content of COX subunits in COX4i1 and COX4i1/COX4i2 KO clones, with COX2 being the most affected. COX4i2 gene KO do not have deleterious effect on COX subunits level. Data in heat map represent log2 values of quantity fold-changes relative to wild-type, according to colour scale shown below the heat map. **d) Representative Western blot analysis of OXPHOS complexes subunits in KO clones.** Whole cell lysates (30 μg of protein) of COX4i1 KO, COX4i2 KO and COX4i1/4i2 KO clones were subjected to SDS-PAGE. Used antibodies are listed in Supplementary Table 1. Actin antibody was used as a loading control. **e) Proteomic analysis of OXPHOS complexes subunits using mass spectrometry label-free quantification (MS LFQ).** Analysis revealed lower levels for majority of cI subunits in COX4i1 and COX4i1/4i2 KO clones. To the contrary, cII, cIII and cV are not significantly affected by COX4I1 gene KO. Data in heat map represent log2 values of quantity fold-changes relative to wild-type. Data in heat map represent log2 values of quantity fold-changes relative to wild-type, according to colour scale shown below the heat map.

In addition to classical electrophoretic and immunodetection approach, relative levels of COX subunits were assessed in a high-throughput manner using mass spectrometry label-free quantification (MS LFQ). Analysis of two technical replicates of two representative clones for each COX4i2 KO, COX4i1 KO and COX4i1/4i2 KO yielded reliable data for eight COX subunits (Fig. 1C). These confirmed our previous findings of generalized COX subunit deficiency in COX4i1 KO and COX4i1/4i2 KO cells, with the most affected subunit being Cox2. COX4i2 KO clones were not significantly changed, as COX subunit amounts were comparable to wild-type cells (Fig. 1C).

Steady-state levels of representative subunits of other OXPHOS complexes (NDUFA9 of cI, SDHA of cII, Core2 of cIII, and F_1_α and Foc of cV) were also studied by SDS-PAGE with Western blot immunodetection (Fig. 1D). Differences between model cell lines were observed in protein level of NDUFA9, which was decreased in COX4i1, COX4i1/4i2, and COX4i2 KO clones, when data were normalized to actin (Fig. S1 C). Upon normalization to citrate synthase, NDUFA9 was significantly, approximately 2-fold lower only in the COX4i1 lacking cells (Fig. S1 D). Core2 subunit was also decreased in COX4i1/4i2 KO, but not in COX4i1 single KO. On the other hand, cII and cV subunits’ levels were similar or even moderately increased in COX4i1/4i2 and COX4i1 KO cells (Fig. S1 D).

The Western blot-based quantification data were complemented by MS LFQ analysis (heat maps in Figure 1E), where quantitative data of 32 cI subunits, all four subunits of cII, seven subunits of cIII, and 13 subunits of cV were obtained. From the perspective of multi-subunit comparison, the decrease of NDUFA9 shown by Western blot analysis was confirmed by LFQ analysis. The levels of cI subunits were between 1.5 – 3-fold decreased in Cox4i1 and COX4i1/4i2 KO clones, while levels of cI subunits in COX4i2 KO were comparable to wild-type. The other OXPHOS complexes (cII, cIII, cV) were not altered in the Cox4i1 and COX4i1/4i2 KO clones. Consistently, COX4i2 KO clones were not significantly affected.

### Compromised OXPHOS function in COX4i1-lacking cells can be complemented by both COX4i1 and COX4i2

To characterize the bioenergetic phenotype of the KO clones, Seahorse Extracellular Flux Analyzer was used to examine rates of mitochondrial respiration and glycolysis (Fig. 2).

**Figure 2:**
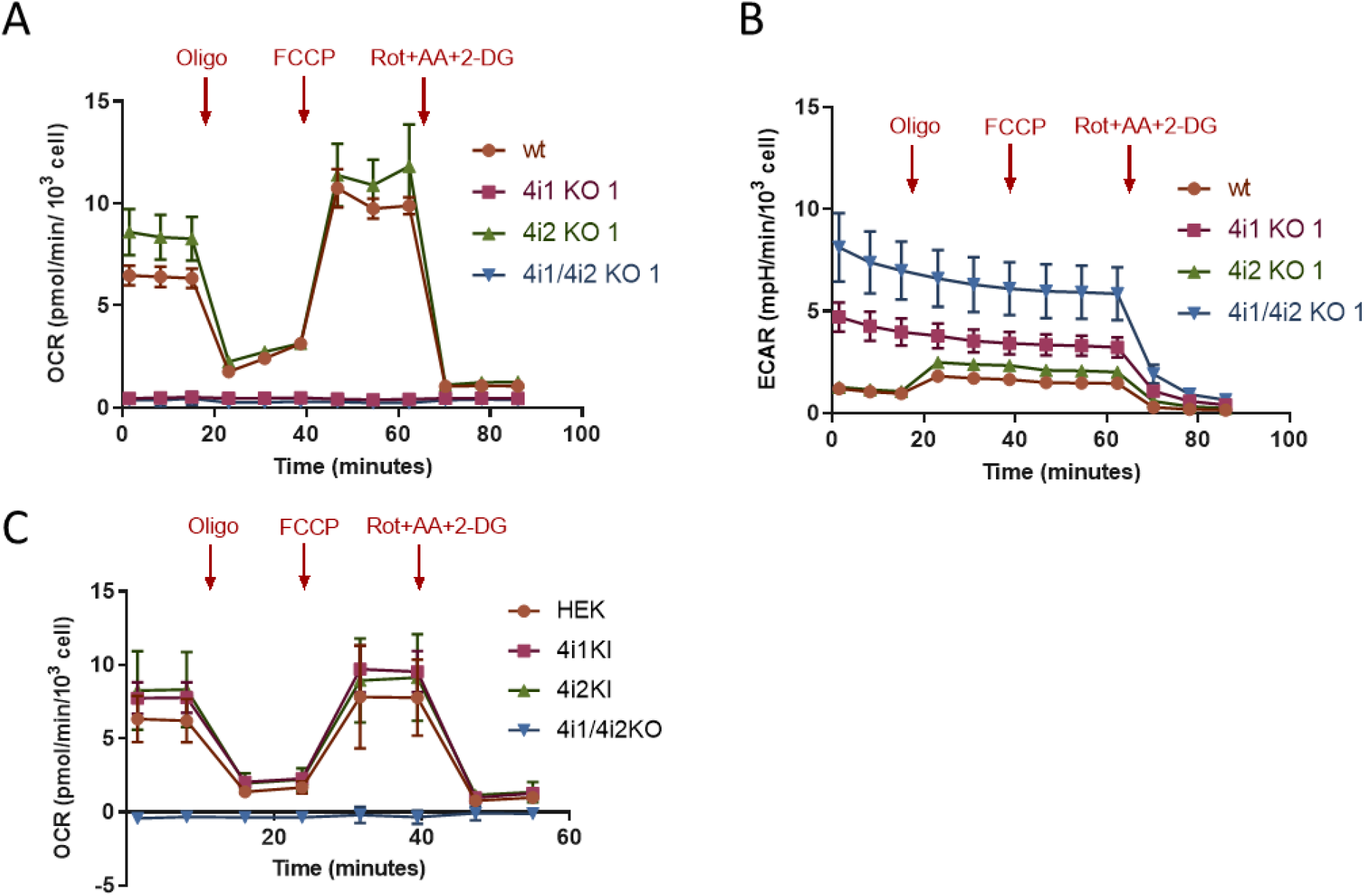
**Representative parallel measurements of a) cellular oxygen consumption rate (OCR) and b) extracellular acidification rate (ECAR)** were performed by Seahorse XF Bioenergetic Analyzer with 10 mM glucose as a substrate, and after subsequent of additions 1 μM oligomycin (Oligo), 1 μM FCCP (FCCP), and inhibitor cocktail of 1 μM rotenone, 1 μg/ml antimycin A, and 100 mM 2-deoxyglucose (Rot+AA+2DG). Data are plotted as mean values ± S.D. of control HEK293 cells (wt, n = 5), COX4i1 KO clone 1 (n = 5), COX4i2 KO clone 1 (n = 4), and COX4i1/4i2 KO clone 1 (n = 3), and demonstrate complete absence of mitochondrial OXPHOS activity in COX4i1 and COX4i1/4i2 KO clones. **c) Complementation of metabolic phenotype in COX4i1 or COX4i2 KI cells.** OCR was recorded by Seahorse XF Bionergetic Analyzer under identical conditions as in a). Data are plotted as mean values ± S.D. of control HEK293 cells (wt, n = 9), COX4i1/4i2 KO (n = 3), COX4i1/4i2 with COX4i1 knock-in (COX4i1 KI, n = 9), and COX4i1/4i2 with COX4i2 knock-in (COX4i2 KI, n = 9).

COX4i2 KO pattern of respiratory and glycolytic fluxes was comparable to wild-type. After recording basal mitochondrial respiration (OCR, oxygen consumption rate, Fig. 2A) and glycolysis (ECAR, extracellular acidification rate, Fig. 2B), addition of oligomycin (inhibitor of cV) caused decrease of OCR, as oxygen consumption coupled to ATP synthesis was inhibited and compensated by increased glycolytic activity represented by ECAR. Addition of chemical protonophore FCCP (which uncouples phosphorylation and oxidation processes) increased OCR, allowing to determine maximal ETC capacity. After last addition, rotenone, antimycin A and 2-deoxyglucose, both OXPHOS and glycolysis were inhibited, therefore OCR and ECAR decreased to minimum and residual background rates were recorded.

As expected, COX4i1 and COX4i1/4i2 KO clones demonstrated complete absence of OXPHOS activity (OCR) or response to additions of uncoupler or inhibitors. Thus, ATP production is independent on OXPHOS. Complete impairment of OXPHOS was compensated by increased glycolytic activity (ECAR) compared to wt cells. Also, ECAR displayed no response to oligomycin, indicating that energy requirements of cells lacking COX4i1 are met by steady rate of glycolysis, likely operating near its full capacity. ECAR rate seemed to be higher for COX4i1/4i2 KO clone than for COX4i1 KO clone. However, it probably reflected unique metabolic settings of individual KO clones.

Importantly, the respiratory OXPHOS impairment was complemented by knock-in (KI) of either COX4i1 or COX4i2 isoforms of COX subunit 4 (Fig. 2C). Expression of either isoform on the COX4i1/4i2 KO background fully restored the respiratory pattern to match control cells.

### Effect of COX4 KO on OXPHOS complexes content and assembly status

To examine effect of OXPHOS subunit changes caused by COX4I1 gene KO, content and assembly status of OXPHOS complexes in native state were studied by BN-PAGE followed by Western blot immunodetection (Fig. 3). Immunodetection with an antibody against COX1 subunit revealed total absence of all assembled forms of COX enzyme in Cox4i1 and COX4i1/4i2 KO clones. Residual COX1 subunit was detected migrating below 100 kDa, likely representing the S1 (early MITRAC) assembly intermediate (Fig. 3A). Early-incorporating subunit COX5a, which also showed residual content in SDS/PAGE experiments, was not detected in association with COX1 in COX4i1 or COX4i1/COX4i2 KO clones (not shown). In contrast, COX4i2 KO displayed a band pattern and amount of COX-containing complexes comparable to wild-type, including COX monomer, dimer, III_2_IV complex and I_1_III_2_CIV_X_ supercomplexes (Figure 3A). cI detection using NDUFA9 antibody in the native gels also confirmed the findings from SDS/PAGE and MS LFQ, showing profound decreased cI content in COX4i1 and COX4i1/4i2 KO clones. Moreover, while in wild-type and COX4i2 KO cI was detected mainly in supercomplexes, the remaining cI in COX4i1-lacking clones was present either as free cI or in association with cIII (I_1_III_2_) (Fig. 3B). cII content was unchanged in COX4i1, COX4i2, and COX4i1/4i2 KO clones compared to wild-type (Fig. 3B). Detection of cIII-containing complexes using antibody against Core2 subunit showed major changes in complex distribution within various assembly forms as a result of COX4i1 absence, rather than changes in enzyme content. While in wild-type and COX4i2 KO cells cIII is detected as dimer (III_2_), III_2_IV and in supercomplexes, lack of COX following COX4i1 gene knock-out shifts almost all cIII to dimer form (Fig. 3C). A mild effect of COX absence was also observed in cV, detected by F_1_β antibody, which shows lower amount of cV dimer, and also minor accumulation of F_1_ and F_1_-c subassemblies in COX4i1 and COX4i1/4i2 KO clones (Fig. 3D). BN-PAGE WB analysis revealed that content of complexes I, III, and IV, as well as their ability to form supercomplexes was restored in cells expressing either of COX4 isoforms on the background of COX4i1/4i2 KO (Fig. 3E). This observation also represents the first direct evidence that COX4i2 can be incorporated into COX complex during *de novo* assembly.

**Figure 3:**
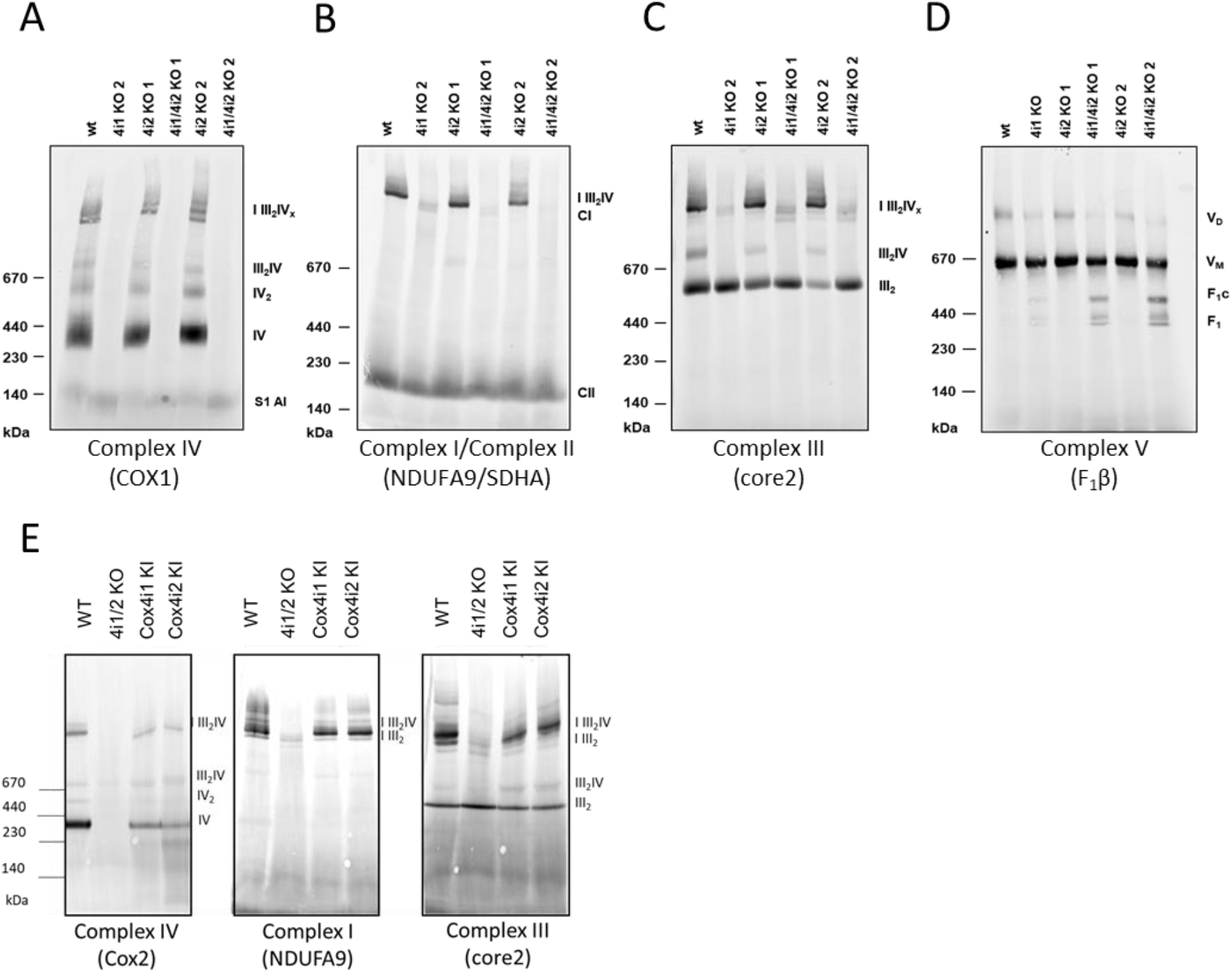
Representative Western blot analysis of OXPHOS complexes and supercomplexes. Digitonin solubilized mitochondria (30 μg of protein) of COX4i1 KO, COX4i2 KO and COX4i1/4i2 KO clones were subjected to BN-PAGE to detect OXPHOS complexes: **a) COX** using COX1 antibody, **b) complexes I and II** using NDUFA9 or SDHA antibody, respectively, **c) complex III** using core2 antibody, **d) complex V** using F_1_β antibody. **e) OXPHOS content and assembly status comparison of COX4i1/4i2 KO and COX4i1 or COX4i2 KI clones.** Digitonin solubilized mitochondria (30 μg of protein) were subjected to BN-PAGE and Western blot detection with indicated antibodies. Used antibodies are listed in Supplementary table 1

Content and assembly status of respiratory chain complexes was further examined in COX4i1/4i2 KO (clone 2) by SILAC/complexome analysis, in comparison to control HEK293 (Fig. 4). Due to the profound COX deficiency in these cells, only two COX subunits, COX5b and NDUFA4, were detected in the KO sample. Both were migrating in low molecular weight region, thus representing unassembled subunits or early assembly intermediates. COX-associated proteins HIGD1A and HIGD2A retained robust signal in KO cells, where a portion of their signal comigrated with MITRAC12 protein. The shift of MITRAC12 migration in KO to the region around 150 kDa in COX4i1/4i2 KO likely represents the COX assembly intermediate that was detected with COX1 antibody in WB experiments, being that MITRAC12 is a COX1-interacting protein [14]. In accordance with BN-PAGE WB analysis and proteomic LFQ studies, SILAC-based complexome profiling confirmed the profound deficiency of complex I in KO cells, its content was approximately five-fold decreased relative to control HEK293. In control cells, complex I was distributed between respirasome (I III_2_IV) and I III_2_ supercomplexes. Interestingly, the low amounts of fully assembled cI in the KO were found either associated to cIII in the I III_2_ supercomplex, or as cI assembly intermediates migrating around 750 kDa. These species contained subunits of all cI assembly modules with the exception of N module components, suggesting it might represent cI precomplex before the addition of the most distal part of peripheral arm harbouring the flavin mononucleotide site. As determined before by WB and immunodetection and LFQ proteomics, only modest decrease of complex III was observed in KO cells. Also in agreement with BN-PAGE WB data, cIII distribution in COX4i1/4i2 KO shifted from supecomplexes to the cIII dimer. Finally, levels of cII were unchanged between control and KO cells.

**Figure 4:**
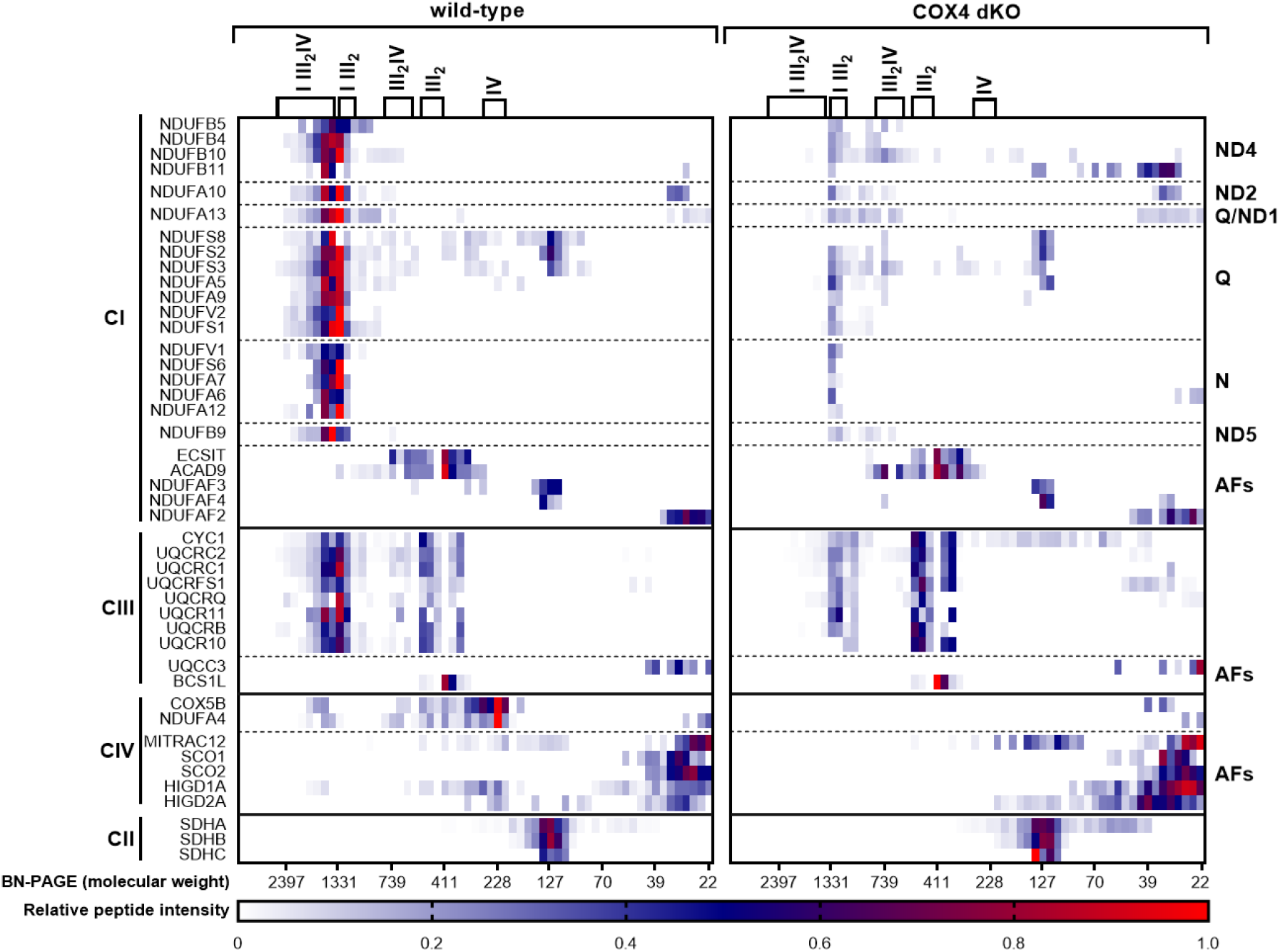
Complexome analysis of OXPHOS complexes content and assembly status. Heatmap displays relative content and migration of respiratory chain complexes components and their assembly factors under native conditions between control HEK293 and COX4i1/4/2 KO cell lines. Data represent relative peptide intensity profile of each protein normalized to the slice with the highest intensity, averaged from reciprocal experiments.

### Decreased rate of mitochondrial proteosynthesis in COX4i1-lacking cells

Pulse-chase *in vivo* ^35^S Met + Cys metabolic labelling of mtDNA-encoded OXPHOS subunits was performed to decipher, whether COX and cI deficiencies detected in electrophoretic analyses were due to subunit degradation following assembly impairment, or rather caused by defect in earlier steps of protein expression. Labelled proteins were separated by SDS-PAGE and detected by phosphor storage screen fluorescence (Fig. 5A). Experiments revealed that majority of mtDNA-encoded OXPHOS subunits, except for ATP6 and ATP8 components of cV, in COX4i1 and COX4i1/4i2 KO clones were synthesized less than in the wt HEK293 cells in the “pulse” samples. On the contrary and consistent with the previous findings, COX4i2 KO clones were not affected, having synthesis rates and protein patterns comparable to wild-type. Detection of labelled proteins following the 24h hour chase in non-labelling medium allowed to estimate protein turnover. In general, levels of labelled proteins were decreasing compared to pulse in all analysed cell lines (Fig. 5A). Strikingly, COX2/COX3 double band completely disappeared during chase in COX4i1 and COX4i1/4i2 KO clones, while residual levels of COX1 were still detected (Fig. 5A, B), reflecting their degradation due to inability to assemble, as suggested by previous data. Concerning other OXPHOS complexes, levels of cIII and cV components displayed higher stability than cI and cIV subunits in cells lacking COX4i1 (Fig. 5A, B), in agreement with data on steady-state levels of these complexes. Therefore, COX deficiency caused by COX4i1 gene KO may have deleterious effect on mitochondrial proteostasis at all levels, affecting the biogenesis of the OXPHOS complexes, especially COX and CI, from synthesis to assembly, as well as their stability.

**Figure 5:**
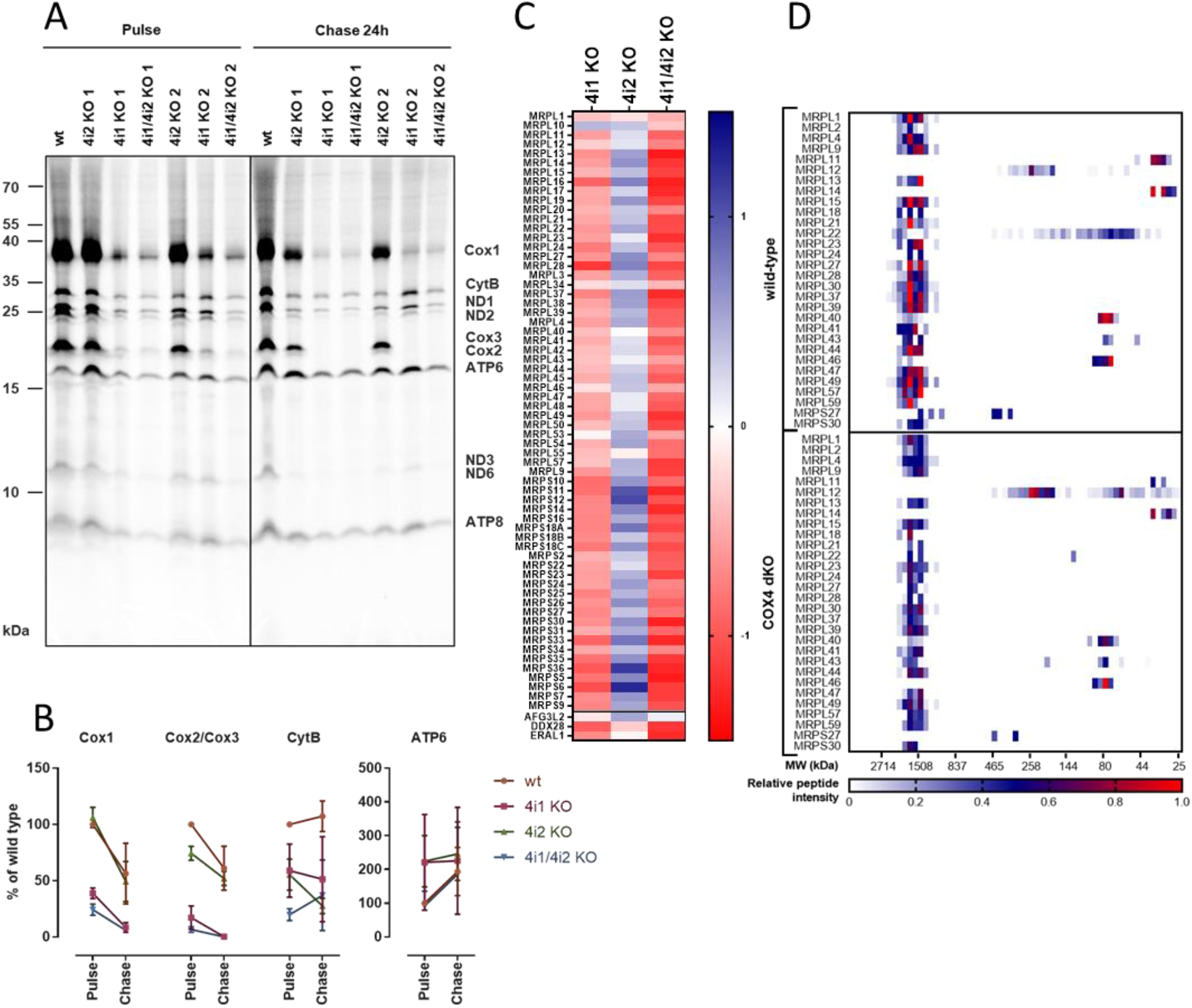
Mitochondrial proteosynthesis a) ^35^S *in vivo* labeling of mtDNA-encoded OXPHOS subunits. Representative image of autoradiographic detection of labelled proteins in 60 μg protein of whole cell lysates analysed by SDS-PAGE, uncovers lower “pulse” level of most mtDNA-encoded proteins due to COX4i1 gene KO. **b) Quantitative analysis of pulse-chase experiments.** Densitometric quantification of labelled COX1, COX2/COX3, CYTB, and ATP6 from autoradiographic images of pulse and 24-hour chase samples are represented as average values of two independent experiments. Error bars represent SD. **c) MS LFQ analysis** reveals lower quantity of mitochondrial ribosomal proteins in COX4i1 and COX4i1/4i2 KO clones. COX4i2 KO clones are comparable to wild-type HEK293 cell line. Data in heat map represent log2 values of quantity fold-changes relative to wild-type. **d) SILAC/complexome profiling of mitochondrial ribosomal proteins.** Heatmap displays relative content and migration of proteins constituting large and small subunit of mitochondrial ribosome. Data represent relative peptide intensity profile of each protein normalized to the slice with the highest intensity, averaged from reciprocal experiments.

Interestingly, the MS LFQ dataset provided further support for the hypothesis of disturbed mitochondrial translation, as content of mitochondrial ribosomal proteins showed a generalized decrease of about 1.5-fold in COX4i1 and COX4i1/4i2 KO clones (Fig. 5C). Also, content of ribosomal assembly factors DDX28 and ERAL1 was 2-fold decreased in COX4i1/lacking cells (Fig. 5C). MS LFQ data were confirmed in the SILAC/complexome profiling experiment, which also showed 1.5-2-fold reduction of both large and small mitoribosome subunits in COX4i1/4i2 KO (clone 2) in comparison with control HEK293 (Figure 5D).

## DISCUSSION

Previously, loss of function (LOF) models of OXPHOS genes in mammalian cells were relying on RNA interference technologies or deriving embryonic fibroblasts from mouse knockout animals (MEFs). The former is complicated with residual content of targeted protein, while generation of MEFs is challenging in terms of resources. Arrival of the relatively straightforward CRISPR-Cas9 gene editing technology [29] thus conveniently broadened experimental repertoire for construction of LOF models in human cell lines.

High-throughput study employing CRISPR Death Screen searching for OXPHOS essential genes in human myelogenous leukemia K562 cells identified COX4i1 as KO-lethal under conditions when galactose was the sole carbohydrate source [37]. With targeted approach, we have prepared single COX4i1 and COX4i2 KO cells, as well as double COX4i1/4i2 KO of both isoforms in HEK293 cells. Even in standard culture conditions, cultivation of COX4i1-lacking clones represented significant challenge due to its highly glycolytic phenotype. Proliferation of KO cells was improved using richer DMEM/F12 medium supplemented with uridine. Therefore, careful selection of a metabolically-flexible cell line and optimization of culturing conditions is necessary for successful experimentation with knock-outs of essential OXPHOS genes.

As COX4 is an early-incorporating subunit [4,10,38], impaired assembly of COX enzyme was expected in COX4i1 and COX4i1/4i2 KO clones. Indeed, native electrophoresis confirmed presence of COX1-containing COX assembly intermediate, most likely representing a MITRAC complex [14]. Fully assembled COX enzyme monomer and dimer were undetected. Moreover, complete absence of fully assembled COX enzyme was coupled with loss of supercomplexes containing COX – III_2_IV and I_1_III_2_IV_1_ (respirasome). Thus, COX4 KO model cell line unambiguously confirmed that COX4 subunit is essential for enzyme assembly process.

Possible compensation of COX4i1 isoform loss by expression of the second isoform, COX4i2, was tested, as this mechanism was described while lost COX7a-H was substituted by its partner, COX7a-L isoform, in COX7a-H KO mice heart [39]. However, Cox4i2 isoform was not detected even after COX4i1 gene KO, thus substitution of missing COX4 isoform 1 by COX4i2 isoform was not activated in HEK293 cells. Therefore, switch of COX4 isoforms seems to strictly depend on oxygen-regulated expression [40,41] and was not triggered by energy crisis in the studied cell line. COX4 isoform switch in HEK293 cells is also improbable since COX4i2 isoform is not expressed in this cell line under normoxia. However, when the expression was forced by transfecting COX4i2 on the COX4i1 KO background, the second isoform was able to complement the COX defect at the structural and functional levels.

Catalytic function of COX enzyme in COX4i1 KO clones was expected to be completely impaired due to loss of catalytic core mtDNA-encoded subunit COX2, indispensable for COX catalytic function [42]. Results of Seahorse measurement confirmed our hypothesis, as COX4i1 KO and COX4i1/4i2 KO clones demonstrated complete absence of oxygen consumption (OCR), thus OXPHOS activity. Impaired OXPHOS function was compensated in COX4i1 KO and COX4i1/4i2 KO clones by increased glycolytic activity (ECAR). Therefore, ATP production is independent on OXPHOS and energy demands of COX4i1 KO cells are covered by ATP production on substrate level, glycolysis. ECAR rate was higher in COX4i1/4i2 KO clone than in COX4i1 KO clone. However, this could be due to individual clonal differences, which originated from single cell selection.

Since COX4i2 is not normally expressed in HEK cells cultured in normoxic conditions, the assembly processes of COX and other OXPHOS complexes (cI, cII, cIII, cV) were not affected by COX4i2 gene KO. Also, COX-containing supercomplexes were detected in COX4i2 KO clones comparably to wild-type. Furthermore, COX4i2 gene KO did not have any impact on mitochondrial respiration, as metabolic phenotype determined by Seahorse was comparable to wild-type. Thus, no significant and functionally relevant changes were observed within COX4i2 KO clones in comparison with wild-type HEK293.

The most affected OXPHOS component apart from CIV in COX4i1 KO and COX4i1/4i2 KO clones was CI, as shown by NDUFA9 immunodetection in BN/PAGE WB and more comprehensively by both label free and SILAC based quantitative proteomics. Previous studies reporting similar cases of COX-cI biogenesis interdependency in mouse cells caused by knock-out of COX10 [25], or by knock-down of COX4 expression [26], concluded that COX absence cause defect in cI assembly. These findings were in line with hypothesis that complexes I, III, and IV need to associate into supercomplexes to successfully proceed through their interdependent assembly pathways [23,24]. Recently, this hypothesis was confirmed by study showing that loss of cIII mtDNA-encoded subunit CYTB impairs assembly of not only cIII but also of cI and cIV Specifically, cI assembly was stalled in late assembly intermediate before addition of matrix-facing N module, which occurs already in pre-respirasome association harbouring cIII and also part of cIV [24]. Accumulation of cI assembly intermediates and the fact that residual mature cI was only found in association with cIII_2_ was also evidenced in our COX4i1/4i2 KO cell line using complexome profiling, indicating that impaired cI assembly is involved in this specific case of cI-deficiency originated by lack of COX assembly. However, this might be only one of the components of the phenomenon as our data also indicate that the interdependency starts already at earlier levels of OXPHOS complexes biogenesis. Indeed, the rate of mitochondrial proteosynthesis was severely reduced both in COX4i1 single or COX4i1/4i2 double knock-out cells. Notably, the synthesis of COX and cI subunits was more affected than cIII cytochrome *b* and mtDNA-encoded subunits of cV. Importantly, such attenuation of mitochondrial translation was not observed in previously published cases of COX-cI interdependency [24–27], indicating that complete absence of COX4 subunit affects COX biogenesis at an earlier step than other COX defects. This might be due to assembly-controlled translational plasticity realized by MITRAC complex, where absence of early-assembly nuclear-encoded subunits such as COX4 leads to arrest of translation of COX1 subunit that remains stalled as a ribosome-nascent chain complex containing MITRAC component C12ORF62 [43]. Therefore, COX enzyme assembly cannot proceed. Moreover, portion of mitochondrial ribosomes might stay occupied by COX1 transcripts, which could explain decreased synthesis of other COX and cI subunits. In cases of interdependency on the basis of COX10 and COX1 mutations, COX4 subunit was detectable and COX1 synthesis was proceeding without translation stalling [25,27]. The COX defect was rather associated with COX1 destabilization due to defective hemylation or C-terminus truncation, respectively. Therefore, another possible link between early phases of cI and COX biogenesis may be represented by MITRAC15 protein. This mammalian homologue of yeast COX1 translational regulator Coa1 was initially identified as COX biogenesis factor [14]. However, it was recently shown that it also serves as a translational factor of cI ND2 subunit, in a similar manner to C12ORF62 function during COX1 synthesis [44]. It could be hypothesized that MITRAC15 could coordinate biogenesis of COX and cI as a bifunctional protein to ensure production of corresponding amount of these respiratory complexes. One could speculate that stalling of COX1 within MITRAC complex may negatively influence the availability of MITRAC15 for ND2 synthesis. ND2 assembly module of complex I serves as a core for addition of other assembly modules [4], therefore its deficiency would have significant effect on cI content. Nevertheless, our data indicate that attenuation of COX and cI mtDNA-encoded subunits is generalized. This could be explained by the decreased content of mitochondrial ribosomal proteins identified by MS LFQ analysis in COX4i1 and COX4i1/4i2 KO cells, as well as by complexome profiling in COX4i1/4i2 KO. The putative mitoribosome insufficiency could be associated with decreased content of ribosomal assembly factors DDX28 and ERAL1 detected in MS LFQ dataset. Contribution of other secondary mechanisms negatively influencing mitoribosome levels, such as energetic insufficiency or changes in redox status in COX4i1-lacking cells should also be considered. However, even residual COX content is sufficient to prevent defect in cI in HEK293, as indicated by study using shRNA-induced knock-down of COX4, which displayed unchanged levels of complex I [45]. Complete clarification of the observed interdependency warrants further studies.

In summary, CRISPR-Cas9 mediated knock-out of COX4 isoforms confirmed its position of early-assembling COX subunit. In normoxia, COX4i2 isoform is not expressed in HEK293, neither is it induced upon removal of isoform 1, therefore COX4i1 single KO and COX4i1/4i2 KO cells displayed same phenotype. Importantly, knock-in of either isoform 1 or 2 into COX4i1/4i2 KO was able to complement COX defect indicating that even COX4i2 is able to initiate *de novo* enzyme assembly. COX4i1-lacking cells presented with COX-cI biogenesis interdependency associated with a yet novel mechanism involving downregulation of mitochondrial protein synthesis most likely by decreasing the content of mitochondrial ribosomes. This reflects the complicated and intertwined mechanisms governing mitochondrial respiratory chain.

## ACKNOWLEDGEMENTS

This work was supported by Grant Agency of the Czech Republic (16-13671S), Grant Agency of Charles University (GA UK 11343/2019), and Ministry of Health of the Czech Republic (NV19-07-00149). Institutional support was provided by Czech Academy of Sciences (RVO:67985823). Project utilized Proteomic core facility, BIOCEV, Faculty of Science, Charles University in Prague (supported by OP VaVpI CZ.1.05/1.1.00/02.0109) for the mass spectrometric measurements.

---

**Supplementary table 1.** List of antibodies used for Western blot detection.

| **Protein specificity** | **Catalog number** | **Provider** | **Dilution** | **Type** |
| --- | --- | --- | --- | --- |
| **COX1** | Ab14705 | Abcam, UK | 1:1000 | mo/M |
| **COX2** | Ab110258 | Abcam, UK | 1:1000 | mo/M |
| **COX3** | Ab110259 | Abcam, UK | 1:250 | mo/M |
| **COX4** | 11463-1-AP | Proteintech Group, USA | 1:1000 | rb/P |
| **COX4i1** | Ab14744 | Abcam, UK | 1:1000 | mo/M |
| **COX4i2** | H00084701-M01 | Abnova, Taiwan | 1:1000 | mo/M |
| **COX5a** | Ab110262 | Abcam, UK | 1:1000 | mo/M |
| **COX6c** | Ab110267 | Abcam, UK | 1:500 | mo/M |
| **Ndufa9** | Ab14713 | Abcam, UK | 1:1000 | mo/M |
| **SDHA** | Ab14715 | Abcam, UK | 1:10000 | mo/M |
| **Core2** | Ab14745 | Abcam, UK | 1:1000 | mo/M |
| **F<sub>1</sub>β** | Ab14730 | Abcam, UK | 1:2000 | mo/M |
| **F<sub>0</sub>c** | Ab18 | Abcam, UK | 1:1000 | rb/P |
| **actin** | MAB1501 | Merck Millipore, USA | 1:60000 | mo/M |
| **Citrate synthetase** | Ab129095 | Abcam, UK | 1:1000 | rb/M |

| **Secondary antibodies** | **Provider** | **Detection wavelength** | **Dilution in TBST** |
| --- | --- | --- | --- |
| **Alexa Fluor 680: goat anti-mouse, donkey anti-goat, goat anti-rabbit IgG** | Thermo Fisher Scientific, USA | 700 nm | 1:3 000 |
| **IRDye 800: goat anti-rabbit, anti-mouse IgG** | Rockland, USA | 800 nm | 1:3 000 |

## Supplementary figures

**Supplementary figure 1.**
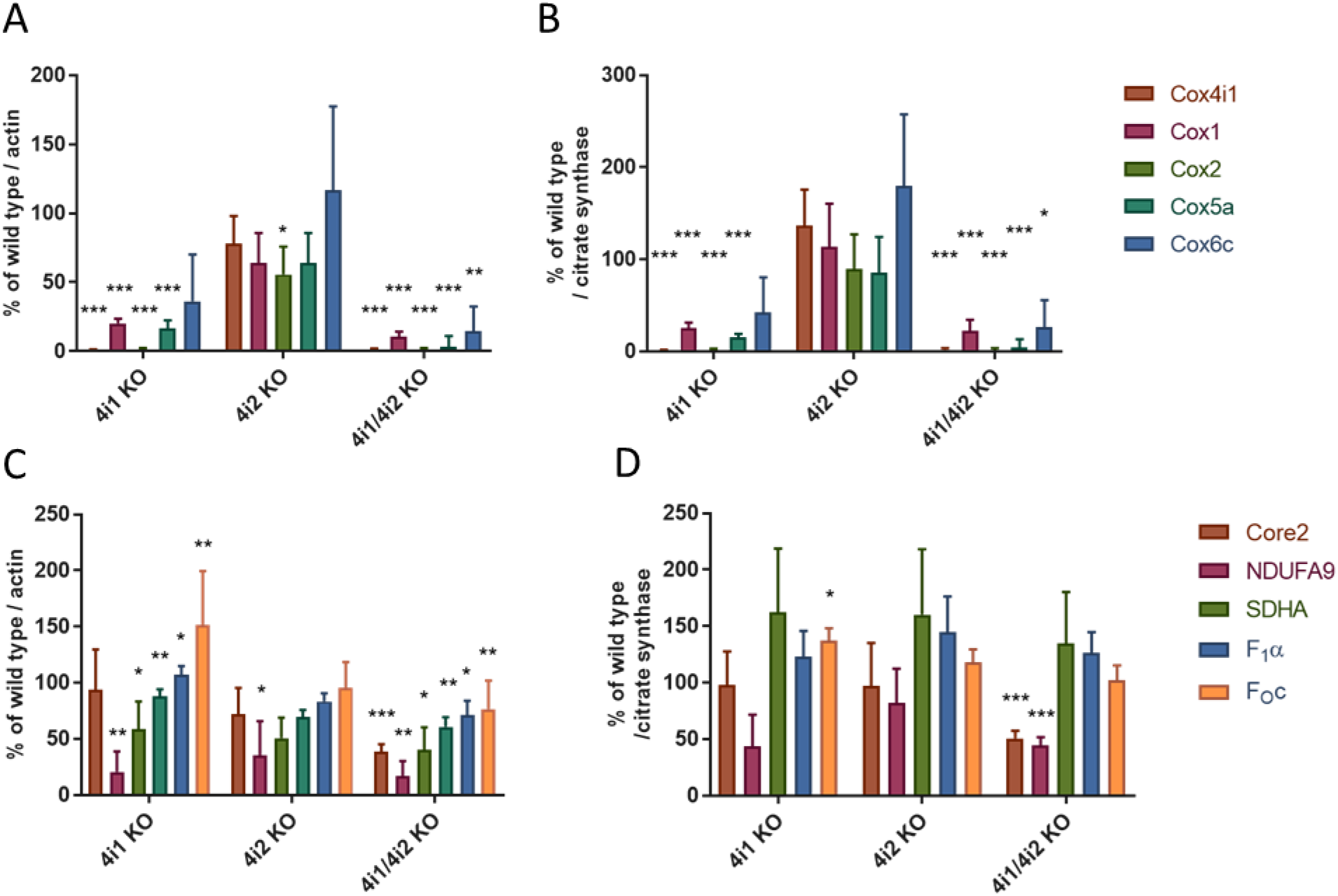
Densitometric quantification of WB experiments a) COX subunits content related to actin. b) COX subunits content related to citrate synthase. c) Quantification of OXPHOS subunits content related to actin. d) Quantification of OXPHOS subunits content related to citrate synthase. Plotted values represent percentage of wild-type. Asterisks (*) represent p-value: * < 0,05; ** < 0,01; *** < 0,001 (related to wild-type). Error bars represent SD. (n = 2)

